# The influence of partner presence and association strength on neophobia in tokay geckos

**DOI:** 10.1101/2023.01.09.523205

**Authors:** Birgit Szabo, Eva Ringler

## Abstract

Neophobia, the avoidance of a novel stimulus, demonstrates the cognitive process of distinguishing novelty from familiarity. Different factors affect neophobia such as evolutionary background, cognitive skills or the social and non-social environment. Individuals from social species often change their neophobic response depending on the presence of a conspecific. Their response might even depend on their social relationship but the directionality of the change varies. To better understand the role of the social environment for neophobia, we tested the effect of the presence/ absence of a mating partner and pair association strength on neophobia across three contexts in the tokay gecko (*Gekko gecko*), a facultative social species with independent offspring. Geckos expressed neophobia similarly when housed singly and in pairs. However, we found that pairs that associated less entered a novel environment faster than those with a strong association while in pair housing. Our study adds important new insights into the relationship between social context and neophobia in a social lizard. Our results highlight that, even in species that express little affiliative behaviour, presence/ absence of a conspecific is insufficient to understand the complex relationship between the social environment and neophobia.

## Introduction

Neophobia is the hesitation to approach or total avoidance of an unfamiliar stimulus due to lack of experience and dissimilarity to known stimuli [1-2]. It is the outcome of the cognitive process that helps individuals distinguish familiar from novel stimuli [1, 3]. On the one hand, such avoidance of novelty can be an ecologically relevant response to evade potentially harmful stimuli, situations, conspecifics or heterospecifics encountered in the environment [1, 3]. On the other hand, avoiding novelty might limit opportunities to discover new resources, to learn or innovate [1, 4-5]. Across species, neophobia differs depending on the trophic level a species occupies [6-8], their evolutionary history (innate recognition) [1, 9-10], their diet [11] and their tendency to exploit new habitats (tolerance to urban environments, invasiveness or frequent migration) [1,12]. While within species, neophobic responses might depend on experience and the ability to generalise across stimuli (cognitive skill) [2], the testing conditions including the familiarity of the environment [1, 13] and the presence or absence of conspecifics [14-20]. Overall, how strongly neophobia is expressed is therefore determined by an individuals’ cognitive skills, the past and current environment (social and non-social) an individual is experiencing and a species’ evolutionary background [1-20].

Encountering novel situations or stimuli can be a stressful experience, especially for social animals when isolated from conspecifics [21-22]. The presence of a conspecific can have a positive effect by acting as a social buffer that reduces stress during an encounter or helps an individual recover faster after the encounter [14, 21-22]. As such, the presence of one or more conspecifics can influence neophobia. Zebra finches (*Taeniopygia guttata*) [15] and budgerigars (*Melopsittacus undulates*) [16], for example, feed earlier from a novel feeder in the presence of conspecifics compared to when alone. Furthermore, group housed dairy calves show less object neophobia than dairy calves in single housing [17]. Contrary, Indian mynahs (*Acridotheres tristis*) take longer to initiate feeding near a novel object when with a conspecific rather than when alone [18] and zebra finches, dogs (*Canis familiaris*), wolves (*Canis lupus*) and ravens (*Corvus corax*) take longer to approach a novel object when with conspecifics compared to when alone [19-20, 23].

Although the mere presence of a conspecific can have an effect on neophobia, their specific relationship with each other should also be considered. For instance, kinship affects approach latency towards novel objects in ravens. Individuals paired with a non-related partner approach novel objects slower than when alone [24]. Male fish (*Oreochromis mossambicus*) show less neophobia in an object neophobia test when they are with a familiar rather than an unfamiliar conspecific or when alone [25]. Finally, in dogs and wolfs, sibling pairs examine novel objects longer compared to non-sibling pairs [19]. These findings highlight that the social context is an important factor mediating neophobic responses in social species [15-20, 23-25]. Depending on which species, test or measure of the social context is considered, however, the directionality (increase or decrease) of neophobic responses changes [15-20, 23-25]. These inconsistencies in results suggest that we still have an incomplete understanding of the significance of different social relationships for neophobia which could even be dependent on the specific social structure of the tested species. Consequently, it is important to consider a broader range of species with diverse social structures to gain a more general understanding of how the social context influences neophobia.

In a previous study [26], we demonstrated that tokay geckos (*Gekko gecko*) show hesitation to attack novel prey items and prey near novel objects. Furthermore, we found that geckos showed the strongest responses towards novel space and demonstrated consistent individual differences in object neophobia. During our first study [26], all geckos were housed singly. Tokay geckos, however, are a facultative social lizard species that shows extended pair association, parental care and temporary family groups, although hatchlings are independent immediately after hatching [27]. Similar to other social species [15-20, 23-25], we hypothesised that geckos might change their neophobic responses depending on the social context they experience during testing. Consequently, we followed up on our first study [26] by retesting the same individuals after being housed in pairs for three months. We predicted that, if a conspecific acted as a social buffer then neophobia would decrease during pair housing compared to single housing. Presence/ absence of a conspecific, however, might be too simple of a measure in this social species and previous studies in other species have shown an effect of the specific social relationship between individuals on neophobia [23-25]. Therefore, we also investigated if the social relationship between the two individuals in a pair might influence neophobia across the three tested contexts. We predicted that pairs with a stronger pair association would show less hesitation to engage with novel stimuli or situations by acting as a more effective social buffer. We define “pair association strength” as frequency of scan sampling points spent in close proximity or touching. Finally, we were interested if we could repeat the findings from our first study [26] comparing latency to attack familiar and unfamiliar prey (food neophobia), latency to attack prey near novel objects (object neophobia) and responses when repeatedly exposed to novel space (space neophobia). In addition, we looked at contextual correlation and individual consistency (repeatability).

## Methods

Our reporting follows the recommendations as laid out by the ARRIVE guidelines [28].

### Animals

We retested the same 22 individuals (10 males: Snout vent length range = 14.005 – 15.796 cm, 12 females: Snout vent length range = 12.493 – 14.339 cm) [27] tested in our first study [26]. While all geckos were housed individually during the first study, in the current study, 18 of them had been housed in pairs for three months (ensuring stable relationships and acclimation to the new housing conditions). All animals were approximately 2-6 years of age at the time of the study and originated from different breeders. We determined the sex based on the presence (male) or absence (female) of femoral glands [27].

### Captive conditions

Nine pairs were housed in large enclosures (90 L x 45 B x 100 H cm) for breeding, while one male and three females were kept singly (male: 90 L x 45 B x 100 H cm; female: 45 L x 45 B x 70 H cm). All enclosures are equipped with a compressed cork wall screwed to the back. We provide refuges (cork branches cut in half, hung on the back wall) as well as cork branches for climbing and life plants as enrichment. Enclosures are set-up bioactive. They contain a drainage layer of expanded clay on the bottom, covered with mosquito mesh (to prevent mixing of the expanded clay and the soil) and topped with organic rainforest soil (Dragon BIO-Ground). Additionally, we spread autoclaved red oak leaves and sphagnum moss on top of the soil to provide shelter and food for the collembola, isopods and earth worms that break down the faecal matter produced by the geckos.

We keep enclosures across two rooms on shelves with small enclosures on the top and large enclosures on the bottom. The environment is fully controlled by an automatic system that aims to mimic the natural conditions geckos experience in nature. Geckos are kept under a reversed 12h:12h photo period (light: 6pm to 6am, dark: 6am to 6pm) with simulated sunrise and sunset which are accompanied by a gradual change in temperature from approximately 25 °C (night cycle) to 31 °C (day cycle). A red light (PHILIPS TL-D 36W/15 RED) not visible to geckos [29] ensures that testing can be done during the night when geckos are active. Additionally, we provide UVB (Exo Terra Reptile UVB 100, 25 W) light from directly above (through the mesh lid of the enclosure) the enclosures during the day cycle. A heat mat (TropicShop) fixed to the outside of each enclosure (increasing the temperature by 4-5 °C) allows lizards to thermoregulation to their optimal body temperature at any time. Base humidity is kept at 50% but daily rainfall every 12h (osmotic water, 30s at 5pm and 4am) increases the humidity to 100% for a short period of time.

### Husbandry

We feed geckos with 3-5 adult house crickets (*Acheta domesticus*) three times per week (Monday, Wednesday, Friday) using 25 cm long forceps. Crickets are gut loaded (cricket mix - reptile planet LDT, Purina Beyond Nature’s Protein™ Adult dry cat food and fresh carrots) to provide optimal nutrition (Vitamin D and calcium). During breeding, we powder crickets with calcium (Exo Terra^®^ Calcium + D_3_) and vitamin powder supplement (Exo Terra^®^ Multi Vitamin) once a week to ensure females have enough calcium to produce eggs. Water is provided *ad libitum* in a water bowl. Lizards’ are weighed once a month and measured (snout vent length) approximately every two months.

### Food and object neophobia

#### Testing procedure

We followed the testing procedure developed in our previous study [26] but describe it here in brief. Geckos were tested in their home enclosures to reduce stress of handling [30] and ensure strong neophobic responses [1,13]. First, we placed a dim white light (LED, SPYLUX^®^ LEDVANCE 3000K, 0.3 W, 17 lm) on top of the tank. Next, we located the focal lizard in the enclosure (removing refuges if necessary). Thereafter, we presented a stimulus in 25 cm long forceps 4-5 cm in front of a geckos’ mouth (optimal to elicit an attack). Geckos were tested once a day (session) every 14 days (inter-session interval). We used different novel foods and objects to be able to test each individual twice. We counterbalanced the order in which lizards were tested: if an individual was first tested on food neophobia it was tested on object neophobia in the second repetition and vice versa. We also randomised testing order within each day to account for order effects. We recoded lizards’ behaviour with a Samsung S20 smartphone (108 Megapixel, 8K-FUHD). Trials were conducted on feeding days (Monday or Wednesday). Testing was done from 6^th^ of April to 16^th^ of May 2022 between 8:00 and 10:15 am.

#### Food neophobia

We used adult house crickets (*Acheta domesticus*) as familiar food. We presented silk moth larvae (*Bombyx mori*) approximately 2.5 cm long and adult cockroaches (*Nauphoeta cinerea*) as the unfamiliar food. New unfamiliar prey was used in each repetition to avoid decreased responses due to repeated exposure (habituation). Geckos had no experience with these prey items for at least one year. Each individual received two presentations per trial in a random order: (1) a familiar cricket and (2) unfamiliar prey. Each presentation lasted for a maximum of 60 seconds.

#### Object neophobia

We used either a course, blue, thin sponge (10 cm L, 2 cm H, 3.8 cm W; Figure 1c) or a fine, blue, high sponge (11.2 cm L, 4.2 cm H, 3.4 cm W; Figure 1d) as novel objects. All novel objects were of similar size and attached to 25 cm forceps. We presented a cricket in forceps as the control condition. Consequently, each gecko received two presentations per trial in a random order: (1) a familiar cricket with no object and (2) familiar cricket next to a novel object. Each object was only used once and we did not interchange objects across individuals. Each presentation lasted for a maximum of 60 seconds.

**Figure 1.**
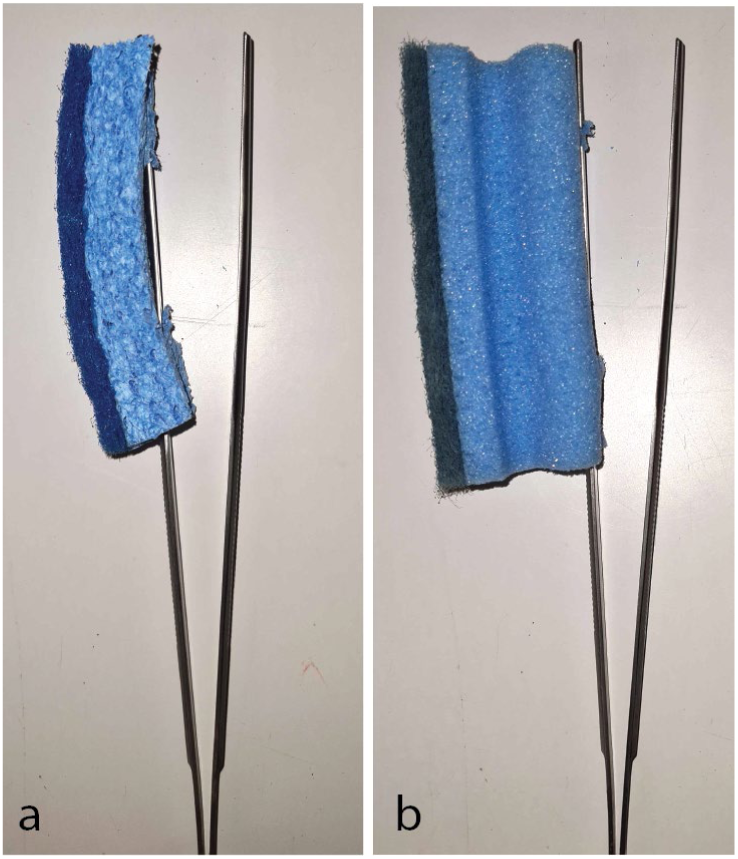
Novel objects used during the object neophobia tests. All objects were attached to 25cm long forceps to be able to present a cricket right next to them during trials. (a) Course, blue, thin sponge (10 cm L, 2 cm H, 3.8 cm W) used in the first trial of object neophobia. (b) Fine, blue, high sponge (11.2 cm L, 4.2 cm H, 3.4 cm W) used in the second trial of object neophobia.

### Space neophobia and exploration

#### Set-up

We followed the general testing procedure developed in our previous study [26] but made some minor changes and describe the changed methods below. Lizards were tested in an empty glass testing tank (45 L x 45 B x 60 H cm, ExoTerra). Four of the sides (three long sides and the bottom) were covered with black plastic on the outside to make them opaque. To be able to measure exploration, we drew a white grid onto the outside of the testing tank (grid: 11.25 cm x 15 cm long sides; 11.25 cm x 11.25 cm lid and bottom; Figure 2b). We placed one testing tank each in an animal room. Testing tanks were placed on top of a table at 100 cm distance facing (with the front transparent doors) a wall. To be able to record videos, we placed a dim white light (LED, SPYLUX^®^ LEDVANCE 3000K, 0.3 W, 17 lm) in the top right corner of the testing tank mesh lid. Trials were recorded from above (40 cm from the tank lid; Figure 2c) using a GoPro (Hero 8; linear mode, 1080 resolution, 24 FPS) mounted on a tripod.

**Figure 2.**
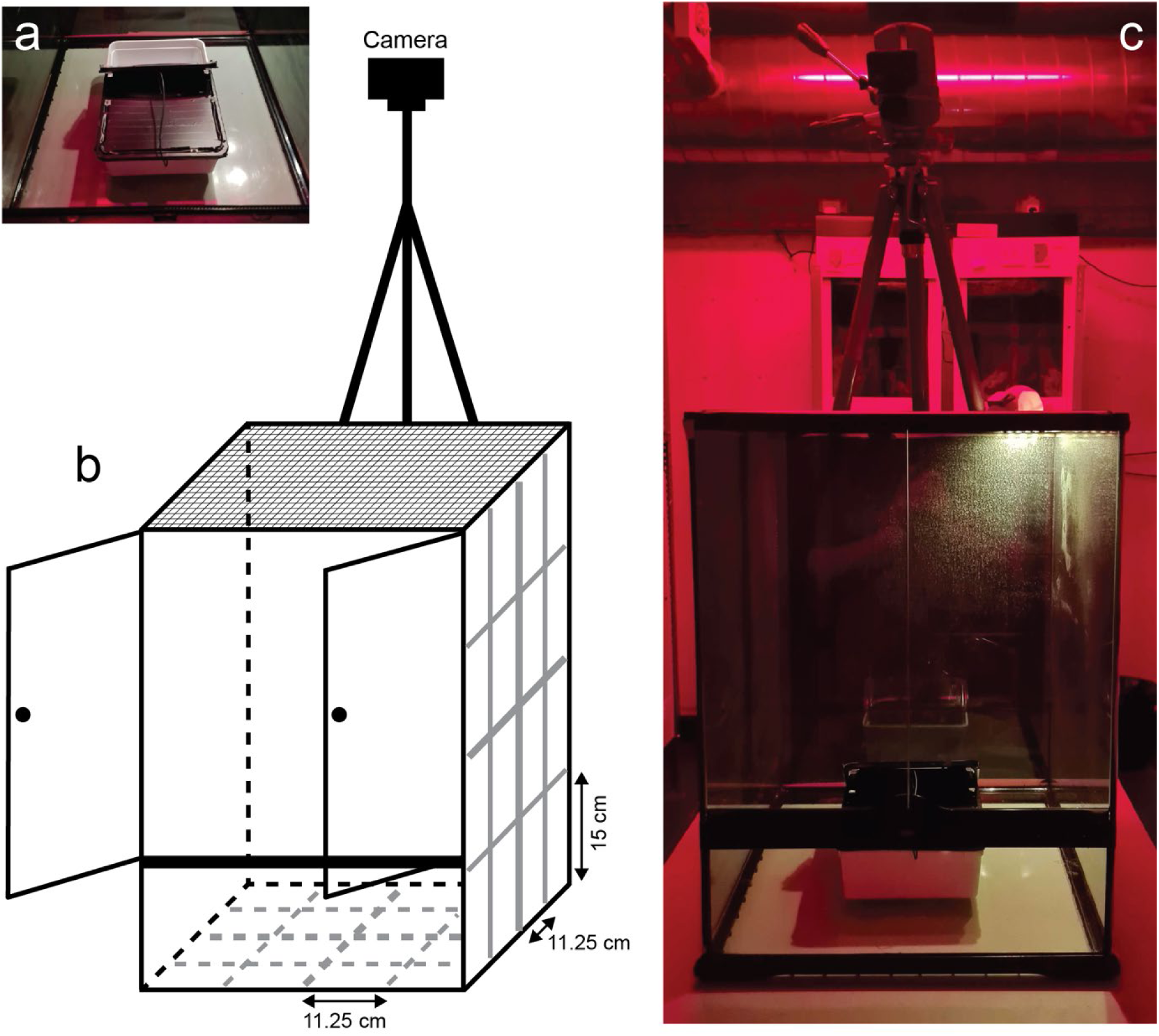
Setup used during the space neophobia test. (a) Picture of the opaque box used to catch lizards (24 cm L x 18 cm W x 7.5 cm H). Picture taken and modified from [26]. (b) Schematic representation of the testing tank (45 L x 45 B x 60 H cm) including the camera. The grid painted on all 6 sides of the testing tank to measure exploration is presented in grey. On the long sides the grid rectangles measured 11.25 cm x 15 cm. On the bottom and the mesh lid, the grid squares measured 11.25 cm x 11.25 cm. (c) Picture of the testing tank including the camera mounted on a tripod and the opaque box inside (grid lines not shown). Sides, except for the front and the lid (made out of mesh), were covered in black plastic to make them opaque. Picture taken from [26].

#### Testing procedure

To test space neophobia, we first captured a focal lizard in an opaque, plastic box (24 cm L x 18 cm W x 7.5 cm H; white opaque bottom with a lid covered in black isolation tape; lids included 6 air holes; Figure 2a). Next, the lizard within the box was carefully placed inside the bottom center of the testing tank with the closed box exit facing the back wall (Figure 2c). Lizards were left to acclimate for 5 minutes. Thereafter, the experimenter started the video recording, opened a third of the box lid carefully (exit), secured the lid to stay open with a wire, closed the testing tank door and left the room. Each individual was left undisturbed for 20 minutes. At the end of the trial, the individual was recaptured either within the opaque box when still inside or using a transparent box that allowing for easier capture. Finally, the lizard was released back into its home enclosure.

To remove chemical cues left by each lizard, the testing tank and opaque box were thoroughly cleaned after each trial with 70% ethanol and left for 15 minutes to allow the alcohol to vanish. Each individual received two trials of space neophobia conducted between 7:45 am and 15:00 pm on the 9^th^ and 23^rd^ of April 2022. Testing took place on non-feeding days (Tuesday and Thursday) with two weeks in between trials.

### Pair association strength

Between 25^th^ of January to 28^th^ of April 2022, we collected data on spatial behaviour within home enclosures using scan sampling. During this time, we recorded the distance between individuals of a pair: (1) more than two snout vent length (SVL) apart, (2) within two SVL, (3) within one SVL or (4) touching. We then used cumulative link mixed models [31] on this ordered categorical data to derive a value for the association strength between paired individuals (based on the pair identity used as a random factor) for each of the nine pairs.

### Data collection

#### Food and object neophobia

We measured capture latency as the time from when a gecko first noticed a prey item (by turning its’ head or eyes) until the first strike regardless of captured success. We scored capture latency using the free behavioural coding software BORIS [32]. To accurately record latency (to 0.001 seconds), we slowed down videos by 50%. If no attack occurred, we recoded occurrence as 0 and assigned this data point a latency of 60 seconds. If the familiar prey/ control prey was not attacked on a given test day, we repeated the whole trial at the end of the experiment. Capture latency of familiar and unfamiliar foods as well as during control and novel object trials recorded during the time when geckos were housed alone were taken from [26].

#### Space neophobia and exploration

We scored the time taken to exit (exit latency, in seconds) into the novel space (testing tank) starting from when the experimenter locked the testing tank door to when a lizard exited the opaque box by lifting its’ tail base over the rim of the box (= exiting with their whole body not counting the tail). If a lizard did not exit the box, we recoded occurrence as 0 and assigned it a latency of 1200 seconds (= 20 minutes). To measure exploration, we counted the number of line crossings after a lizard had exited the box. If a lizard crossed in a corner, we counted two line crossings. To accurately estimate each individuals’ exploration score we divided the total number of line crossings by the time left for exploring after the opaque box was exited. Exit latencies recorded during the time when geckos were housed alone were taken from [26].

### Inter-observer reliability

Due to the nature of the study, we were unable to score behaviour blind as to test (food, object or space neophobia) and stimulus (familiar/ control, unfamiliar). Therefore, 40% of videos were scored by an observer that was unaware of the objectives of the study. Latency and exploration measures between the blind observer and the original observer (BS) were highly consistent (Latency: Spearman rank correlation, r_s_ = 0.973; exploration: Spearman rank correlation, r_s_ = 0.986).

### Statistical analyses

#### Changes in neophobia from single to pair housing

To investigate if neophobic responses changed from single to pair housing, we only focused on data from those individuals that were housed in pairs in the current study (excluding 3 females and 1 male). We ran Bayesian Generalised Linear Mixed Model (GLMM, package MCMCglmm [33]) with a Gaussian family for each test (food, object and space neophobia) using the log-transformed latency towards the unfamiliar/novel stimulus as the response variable. We log-transformed latencies to better fit a normal distribution. In all three models, we included study and test trial (both studies combined, 1-4 trials depending on the test) as fixed effects. In the model looking at food neophobia, we included temperature as an additional fixed effect while we included individual testing order in the model looking at space neophobia. We also calculated adjusted repeatability across both studies accounting for stimulus (food and object neophobia) or test trial (space neophobia) (package rptR [34]).

#### Changes in neophobia associated with pair association strength

To investigate if the strength of the association between individuals within a pair was correlated with neophobia (food, object and space) and exploration, we ran Bayesian GLMMs (one for each test) with a reduced dataset including only those individuals for which we had data on pair association strength. Again, log-transformed latencies/ exploration score were used as the response variable in all models. All models included a fixed effect of trial in interaction with association strength. In the models focusing on food and object neophobia we included stimulus in interaction with association strength as another fixed effect. Additionally, in the model focusing on food neophobia we also included temperature as a fixed effect, while in the model focusing on space neophobia we included testing order. In all models, trial was used as a categorical variable to represent repetitions in time. We investigated significant results of interactions using PostHoc tests for estimated marginal means of linear trends (EMM, package emmeans [35]).

#### Neophobia, repeatability and contextual correlation

Similar to our first study [26], we investigated if neophobia occurred in the food, object and space neophobia test and if behaviour was repeatable. To this end, we ran a Bayesian GLMM for each test with the log-transformed latency as the response variable on the data of all 22 individuals. As fixed effects we included (1) stimulus (familiar/ control, unfamiliar/ novel) to investigate if neophobia occurred (food and object neophobia only), (2) trial to investigate differences across stimuli, (3) sex (male or female), (4) room (room 2 or room 5), (5) average enclosure temperature, (6) individual testing order, and (7) body condition (scaled mass index [36]). The same fixed effects were used to investigate exploration with the log-transformed exploration score as the response variable. We also calculated adjusted repeatability within tests accounting for stimulus (food and object neophobia) or trial (space neophobia).

Across studies, we noticed differences in responses depending on the type of unfamiliar food presented. To look into this in more detail, we ran a Bayesian GLMM with a Gaussian family using the log-transformed capture latency (from both studies) as the response variable and food type (cricket, locust, mealworm, silk moth larvae or cockroach) as the fixed effect. We then calculated the difference in responses across food types using the posterior produced by the model.

To compare neophobic responses across tests, we only focused on latency in response to unfamiliar/ novel stimuli. To make latency comparable across tests, we calculated relative latencies by dividing capture latency in the food and object neophobia test by 60 (maximum trial time in seconds) and the exit latency in the space neophobia test by 1200 (maximum trial time in seconds). We then ran a Bayesian GLMM to investigate differences in relative latency (log-transformed response variable) across tests (fixed effect). Additionally, we calculated average latencies shown towards unfamiliar stimuli across trials and used spearman rank correlation tests to investigate if latencies correlated across tests. Finally, we calculated adjusted repeatability across tests accounting for stimulus and test.

All analyses were run in R version 4.0.3 [37]. We report results as *p* > 0.1 no evidence, 0.1 < *p* < 0.05 weak evidence, 0.05 < *p* < 0.01 moderate evidence, 0.01 < *p* < 0.001 strong evidence, *p* < 0.001 very strong evidence [38].

Models investigating food/ object neophobia and differences across contexts included a random intercept of animal identity and a random slope of trial nested in session. Models investigating space neophobia and exploration included a random intercept of animal identity and a random slope of trial. We included a second random effect in models looking at the effect of pair association strength with an intercept of tank number and a random slope of trial nested in session. Random effects account for repeated testing, autocorrelation across successive samples and that the male and the female in a pair always received the same value of association strength. For all Bayesian models, we made sure that no autocorrelation occurred (correlation between lags < 0.1 [33]), that the MCMC chain sufficiently mixed (by visually inspecting plots [33]) and was run long enough (Heidelberg and Welch diagnostic tests [33]).

### Ethical statement

Our tests were strictly non-invasive and followed the guidelines provided by the Association for the Study of Animal Behaviour/ Animal Behaviour Society for the treatment of animals in behavioural research and Teaching [39] and the Guidelines for the ethical use of animals in applied animal behaviour research by the International Society for Applied Ethology [40]. Experiments were approved by the Suisse Federal Food Safety and Veterinary Office (National No. 33232, Cantonal No. BE144/2020). Captive conditions were approved by the Suisse Federal Food Safety and Veterinary Office (Laboratory animal husbandry license: No. BE4/11).

## Results

### Changes in neophobia from single to pair housing

We found very strong evidence that neophobic responses towards unfamiliar food increased from single to pair housing (GLMM, estimate = 4.388, CI_low_ = 2.334, CI_up_ = 6.205, p = 0.00007; Figure 3), that the latency to attack unfamiliar prey decreased across test trials (GLMM, estimate = -1.176, CI_low_ = -1.933, CI_up_ = -0.410, p = 0.003) and that neophobic responses also decreased with decreasing temperature (GLMM, estimate = -1.480, CI_low_ = -2.443, CI_up_ = - 0.518, p = 0.003; supplementary material Table S1). We found no evidence that neophobic responses towards novel objects changed from single to pair housing (GLMM, estimate = 0.367, CI_low_ = -0.638, CI_up_ = 1.435, p = 0.485; Figure 3) or that test trial was related to latency (GLMM, estimate = -0.283, CI_low_ = -0.765, CI_up_ = 0.204, p = 0.244; supplementary material Table S2). We found no evidence that neophobic responses towards novel space changed from single to pair housing (GLMM, estimate = 0.503, CI_low_ = -0.100, CI_up_ = 1.113, p = 0.107; Figure 3) but found strong evidence that space neophobia decreased with test trial (GLMM, estimate = -0.627, CI_low_ = -0.982, CI_up_ = -0.259, p = 0.002) and no evidence that space neophobia was affected by individual testing order across studies (GLMM, estimate = -0.025, CI_low_ = -0.075, CI_up_ = 0.026, p = 0.340; supplementary material Table S3). Overall, we found that individual responses were only repeatable in the object neophobia test (R = 0.124, Ci_low_ = 0.006, CI_up_ = 0.254, p = 0.008) but not in the food (R = 0.004, CI_low_ = 0, CI_up_ = 0.098, p = 1) or space neophobia test (R = 0.186, CI_low_ = 0, CI_up_ = 0.458, p = 0.090).

**Figure 3.**
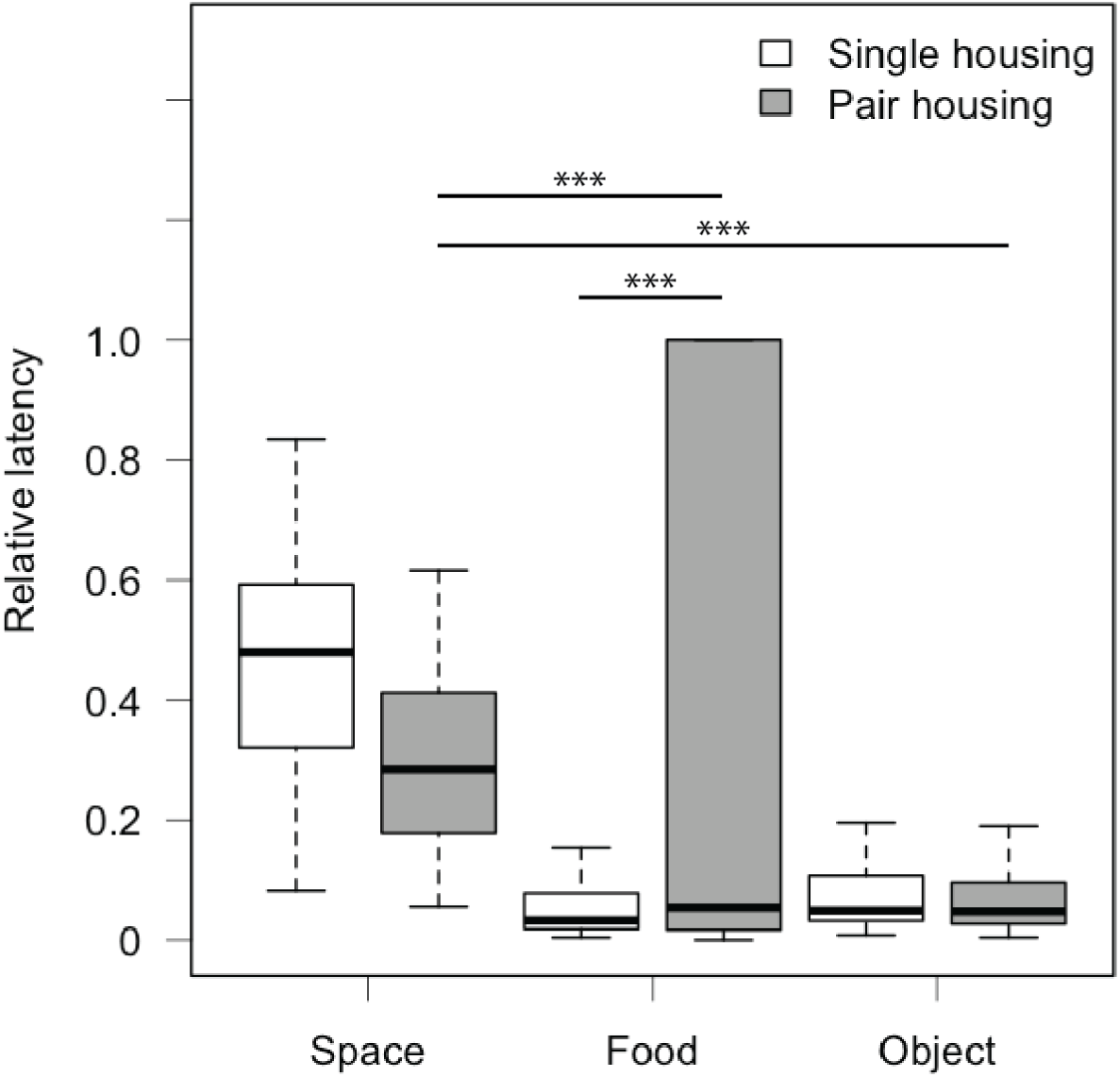
Relative latency shown towards unfamiliar/ novel stimuli across tests (space, food and object neophobia) and studies (single and pair housing study). The bold line within boxes indicates the median, the upper box edges give the upper quartile, the lower box edges the lower quartile, the top whisker ends the maximum and the bottom whisker ends the minimum (outliers are not shown). *** p < 0.001.

### Changes in neophobia associated with pair association strength

In the food neophobia test, we found no evidence for a correlation of pair association strength with the latency to attack familiar or unfamiliar food (GLMM, estimate = 0.279, CI_low_ = -1.901, CI_up_ = 2.306, p = 0.792), or across trials (GLMM, estimate = 1.453, CI_low_ = -1.369, CI_up_ = 4.210, p = 0.294; supplementary material Table S4). Similarly, we found no evidence for a correlation of pair association strength with the latency to attack control crickets or crickets near novel objects (GLMM, estimate = 1.182, CI_low_ = -0.488, CI_up_ = 2.828, p = 0.158), or across trials (GLMM, estimate = -0.869, CI_low_ = -3.413, CI_up_ = 1.618, p = 0.484; supplementary material Table S5) in the object neophobia test. In the space neophobia test, we found moderate evidence for a correlation of association strength with the latency to exit into a novel environment across trials (GLMM, estimate = 1.050, CI_low_ = 0.109, CI_up_ = 1.942, p = 0.027; supplementary material Table S6). PostHoc estimated marginal means of linear trends showed that pairs with a weaker association exited faster into the novel environment in trial 2 (EMM: trend = 1.150, CI_low_ = 0.389, CI_up_ = 1.904; Figure 4, supplementary material Table S6). Finally, we found no evidence for a correlation of pair association strength with exploration across trials (GLMM, estimate = -0.763, CI_low_ = -2.081, CI_up_ = 0.507, p = 0.238; supplementary material Table S7).

**Figure 4.**
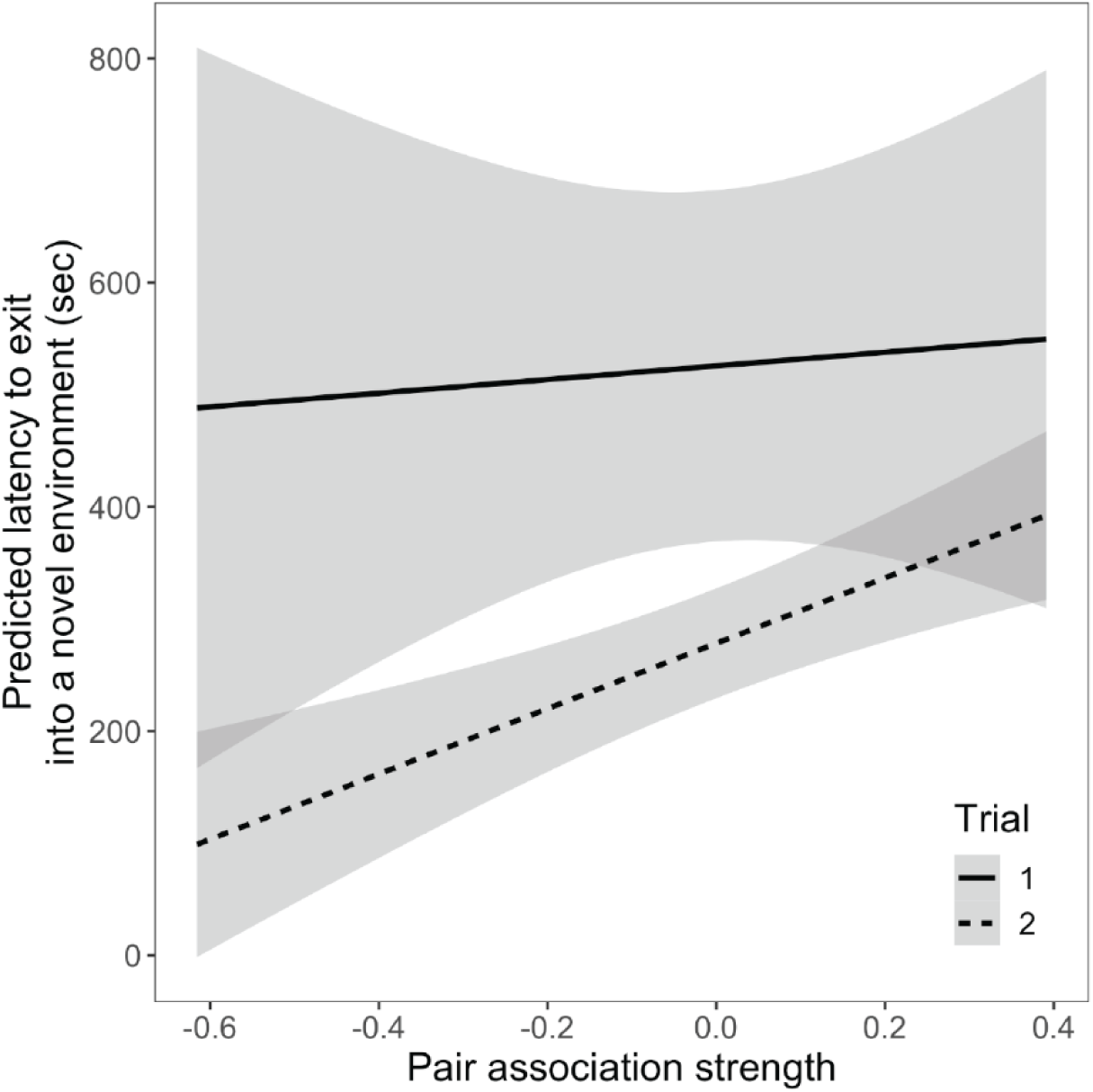
Relationship between the latency to exit into a novel space and pair association strength. Data are separated into trial 1 (solid line) and trial 2 (dashed line) of the space neophobia test. The shaded area represents the 95% confidence interval.

### Neophobia, repeatability and contextual correlation

#### Food neophobia

We found strong evidence that lizards attacked unfamiliar prey slower compared to familiar crickets (GLMM, estimate = 1.021, CI_low_ = 0.375, CI_up_ = 1.653, p = 0.003; Figure 5a). We also found strong evidence that lizards attacked cockroaches faster than silk moth larvae (GLMM, estimate = -1.031, CI_low_ = -1.794, CI_up_ = -0.319, p = 0.003). Additionally, we found strong evidence that lizards attack latency decreased with rising temperatures (GLMM, estimate = - 1.497, CI_low_ = -2.484, CI_up_ = -0.579, p = 0.004). We found no evidence that the other fixed effects affected attack latency (supplementary material Table S8). When looking at the different food types across both studies, we found evidence that lizards took the longest to attack silk moth larvae compared to all other food types (GLMM, supplementary material Table S9; Figure 6). They also took longer to attack mealworms (GLMM, estimate = 0.940, CI_low_ = 0.170, CI_up_ = 1.701; Figure 6) and locusts (GLMM, estimate = 0.776, CI_low_ = 0.025, CI_up_ = 1.521; Figure 3) compared to cockroaches. Individuals’ attack latency was not repeatable within the food neophobia test (R = 0, CI_low_ = 0, CI_up_ = 0.197, p = 1; Figure S1).

**Figure 5.**
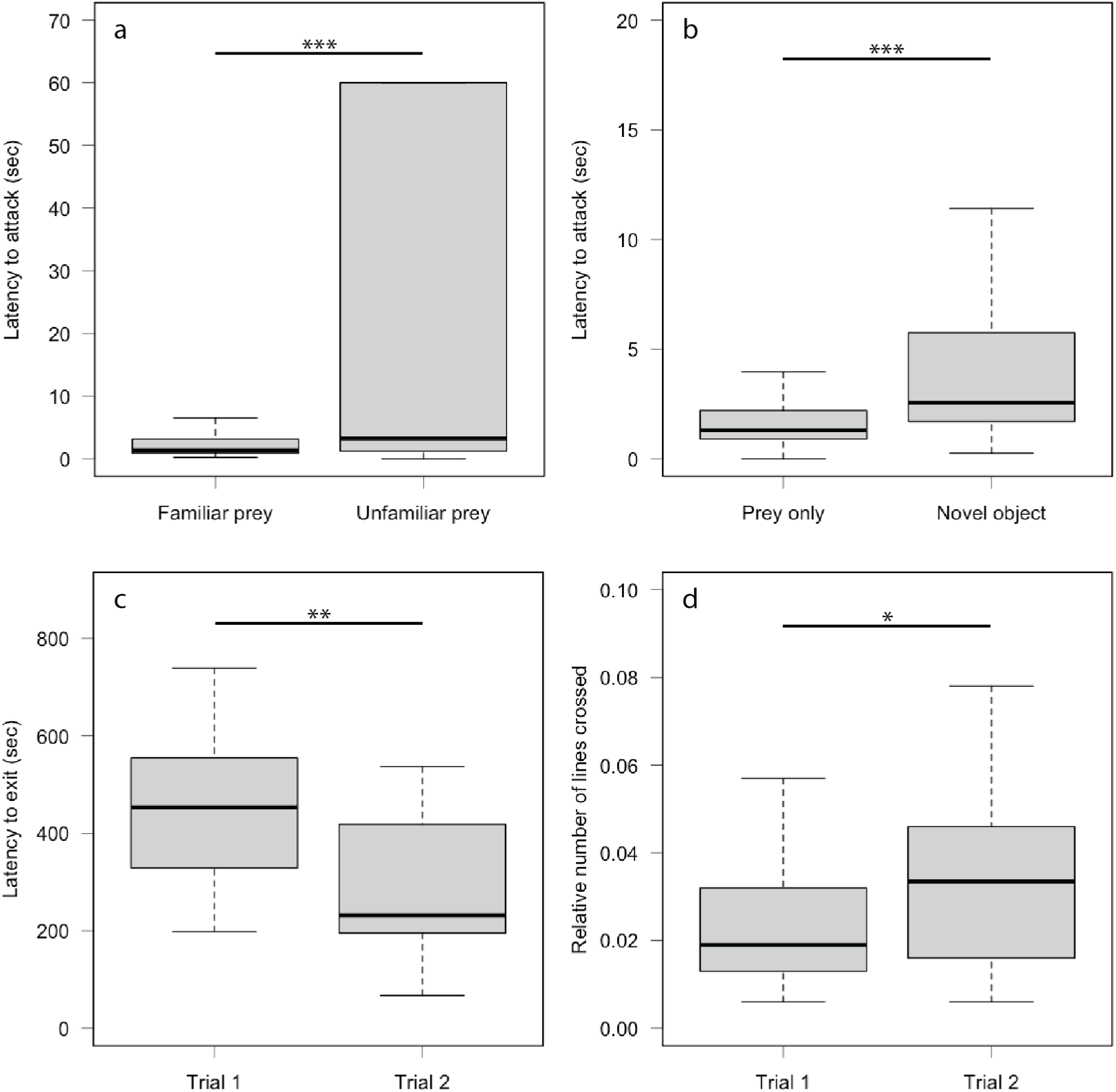
Boxplots of results obtained in the food, object and space neophobia test. (a) Attack latency towards familiar (cricket) and unfamiliar (response towards silk moth larvae and cockroaches pooled) prey. (b) Attack latency towards familiar prey alone (control) and towards familiar prey near novel objects (both sponges pooled). (c) Latency to exit into the novel environment in trial 1 and 2. (d) Exploration score in trial 1 and 2 of the space neophobia test. The bold line within boxes indicates the median, the upper box edges give the upper quartile, the lower box edges the lower quartile, the top whisker ends the maximum and the bottom whisker ends the minimum (outliers are not shown). * p < 0.05, ** p < 0.01, *** p < 0.001.

**Figure 6.**
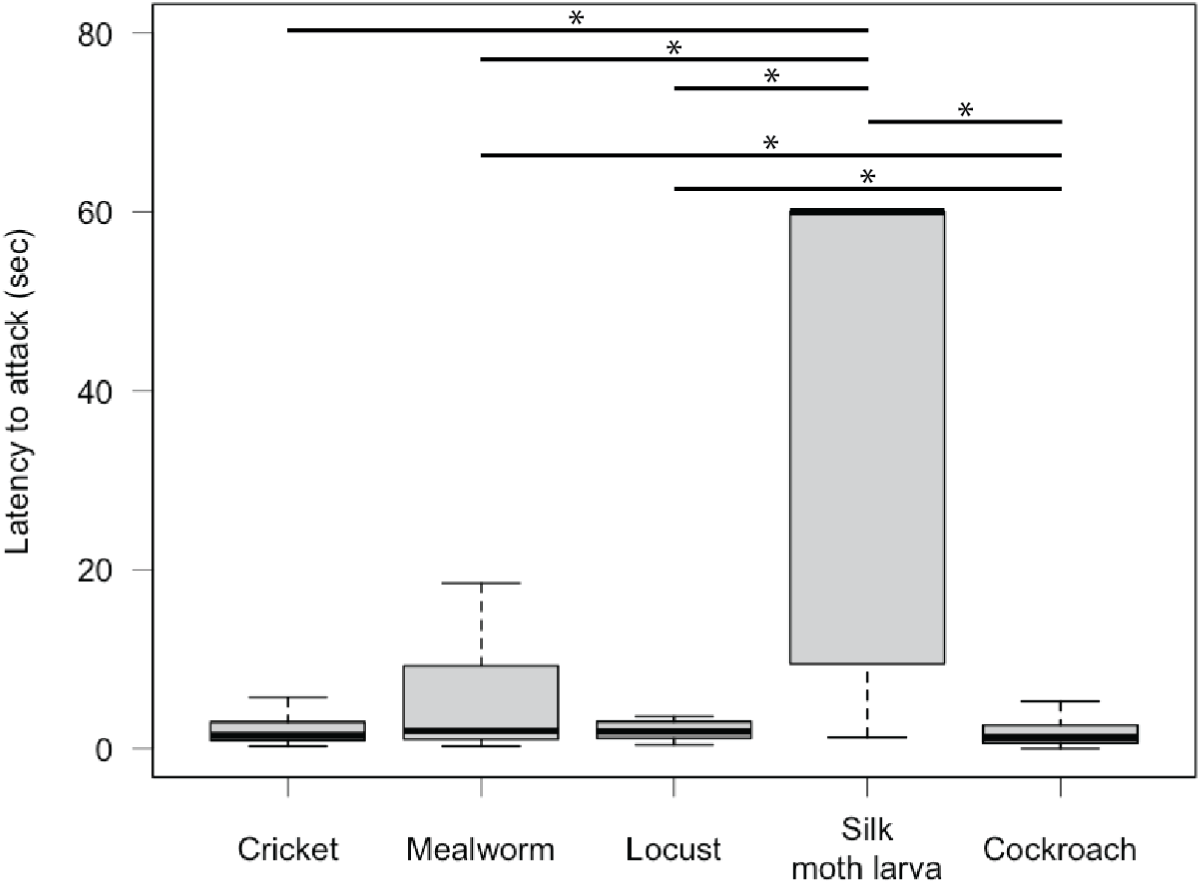
Box plots summarising capture latencies in response to the different prey items used across both studies. The bold line within boxes indicates the median, the upper box edges give the upper quartile, the lower box edges the lower quartile, the top whisker ends the maximum and the bottom whisker ends the minimum (outliers are not shown). * p < 0.05.

#### Object neophobia

We found strong evidence that lizards attacked familiar prey near novel objects slower compared to the control cricket (GLMM, estimate = 0.992, CI_low_ = 0.462, CI_up_ = 1.517, p = 0.003; Figure 5b). We found no evidence that the other fixed effects affected attack latency (supplementary material Table S10). We found moderate evidence that individuals’ attack latency was not repeatable within the object neophobia test (R = 0.225, CI_low_ = 0, CI_up_ = 0.448, p = 0.013; Figure S1).

#### Space neophobia and exploration

We found very strong evidence that exit latency decreased from trial 1 to trial 2 (GLMM, estimate = -0.491, CI_low_ = -0.821, CI_up_ = -0.155, p = 0.006; Figure 5c). We found moderate evidence that exit latency decreased the later in the day lizards were tested (GLMM, estimate = -0.109, CI_low_ = -0.203, CI_up_ = -0.022, p = 0.023) and we found weak evidence that lizards with lower body condition also took longer to exit into the novel environment (GLMM, estimate = -0.012, CI_low_ = -0.026, CI_up_ = 0.001, p = 0.071). We found no evidence that the other fixed effects affected exit latency (supplementary material Table S11). We found no evidence that individuals’ exit latency was repeatable (R = 0.044, CI_low_ = 0, CI_up_ = 0.468, p = 0.418; Figure S1).

We found moderate evidence that exploration increased from trial 1 to trial 2 (GLMM, estimate = 0.459, CI_low_ = 0.010, CI_up_ = 0.896, p = 0.044; Figure 5d, supplementary material Table S12). We found no evidence that the other fixed effects affected exploration scores (supplementary material Table S12). We found strong evidence that individuals’ exploration score was repeatable (R = 0.538, CI_low_ = 0.152, CI_up_ = 0.781, p = 0.004; Figure S1).

#### Contextual correlation

We found strong evidence that lizards showed the strongest responses towards exiting into a novel environment compared to attacking unfamiliar prey (GLMM, estimate = 1.336, CI_low_ = 0.729, CI_up_ = 1.977, p = 0.00007) or attacking prey near novel objects (GLMM, estimate = - 1.539, CI_low_ = -2.117, CI_up_ = -0.900; Figure 3, supplementary material Table S13). We found no evidence that responses towards novel stimuli were correlated across tests (supplementary material Table S14). Finally, we found no evidence that individual neophobic responses were repeatable across tests (R = 0.037, CI_low_ = 0, CI_up_ = 0.134, p = 0.167).

## Discussion

Overall, the presence of a second individual had no effect on object and space neophobia per se, but we found pair association strength to significantly impact space neophobia. Individuals from pairs with a weaker association exited faster into a novel environment. Food neophobia increased during pair housing but rather than social context, prey type is more likely to explain this change. Our follow-up study confirmed the results obtained from the first study [26]. Geckos hesitate to attack unfamiliar food, familiar food near novel objects and when entering novel space but showed the strongest responses towards novel space. Furthermore, individual repeatability was low within test as well as across tests. We only found high repeatability in our measure of exploration in novel space. Contrary to the first study, we found no correlation across contexts. Finally, across studies we found no repeatability except for object neophobia which was low and (expected) habituation in response to repeated exposure to the same novel space.

The main aim of this study was to investigate the effect of a social partner and their relationship on neophobia. The mere presence of a second individual did not affect responses towards familiar or novel stimuli showing no social buffering. While the relationship between individuals within a pair affected space neophobia but not food and object neophobia or exploration. When entering a novel space, individuals with a weaker association with their partner entered the novel environment faster compared to individuals that had a stronger association. This effect was strongest in the second space neophobia trial, although we found a similar positive relationship in the first trial. In the wild, male tokay geckos establish territories from which they call during the breeding season to attract females [27]. They form pairs and later, after the offspring hatch, temporary family groups until offspring become sexually mature and are evicted from the territory [27]. It could be that those individuals that do not bond well with their chosen partner are less averse to novel space possibly to move on and find another better suited mate. Previous work has demonstrated lower neophobia in animals when migrating possibly because they encounter novelty regularly during their journey [1, 41]. Furthermore, pair bond stability is related to reproductive success [42] and mate familiarity leads to earlier reproduction in at least one lizard species [43]. It could, therefore, be an adaptive strategy to find the most suitable mate and produce the highest number of offspring within a breeding season by exploring outside the familiar home range. Based on tokay gecko ecology, we would actually expect to find a stronger effect in females rather than males because females are the moving and choosing sex. However, our low sample size of only nine mating pairs precluded a sex separated analysis. It will be interesting to repeat this study with additional pairs in the future, to investigate if there is a difference in males and females regarding the effect of pair association strength on space neophobia.

The results from this study confirmed the overall results of our first study, when lizards were still housed singly [26]. In both studies, geckos hesitated to attack unfamiliar prey in comparison to familiar prey, to attack familiar prey near novel objects compared to prey alone and to enter a novel space. Furthermore, in both studies geckos hesitated the most when entering a novel space. Although this might be caused by generally higher space neophobia in this species, it could also reflect differences in motivation across tests. Compared to the food and object neophobia tests, the space neophobia test did not present food and lower motivation could therefore have led to increased hesitation to engage with novelty. Across studies, we also found differences in attack latency across novel foods. Based on our results it seems that worm shaped prey elicits the strongest hesitation possibly because geckos main prey in the wild constitute of moths, crickets and grasshoppers [44]; these food types were also attacked the fastest in our test. This could point towards an innate preference similar to what has been shown in other species (e.g. innate prey preference [45]; innate colour preference [46]) but should be confirmed by testing individuals that have never encountered grasshoppers or cockroaches before. Consequently, the change in responses towards novel food from the first to the second study seem more likely caused by prey type rather than changes in the social environment. Finally, contrary to the first study, we did not find a correlation between contexts. We could not reproduce this result in the current study possibly confirming our previous conclusion that this correlation could be an artefact of the study. Nevertheless, the previously found correlation might have been masked due to geckos’ preference for cockroaches and aversion to consume silk moth larvae combined with the effect of habituation to the novel environment. Consequently, to confirm the absence of a correlation we would need to test a cohort of naïve individuals.

Similar to our previous results [26], we found no or very low individual repeatability within tests, across tests and even across studies. Previously, we found low individual repeatability in the food neophobia test (R = 0.101), and in the current study, we found no repeatability (R = 0) likely caused by the huge difference in response to the novel food items used. More than half of all tested individuals did not attack silk moth larvae while all individuals attacked cockroaches sometimes even faster than control crickets (Figure S1a). The strong influence of prey species on responses is further confirmed by the low repeatability (R = 0.004) when all trials across both studies are taken into account. Our food neophobia test, therefore, seems unreliable in tokay geckos due to the strong effect of prey species. In the first study, we found individual repeatability in object neophobia (R = 0.190) [26], which we could confirm with new novel objects (R = 0.225) albeit not significantly. Object neophobia was the only trait that was also repeatable across studies (R = 0.124) making it a reliable measure for future investigations in this species. However, individual repeatability was still low which might have been caused by the low number of repetitions (N = 4). In the space neophobia test we found no individual repeatability for exit latency but quite high individual repeatability for exploration (R = 0.538). Habituation had a strong effect on exit latency while it only had a weak effect on exploration possibly leading to the differences across these two measures. Rather than just exit latency, the decrease in exit latency (degree of habituation) during the repeated exposure to the same novel space could be an interesting measure to consider in future studies. The individual repeatability we found for exploration, generally, falls well within the range reported for behaviour [47-48] and demonstrates that our test methodology is well suited to measure exploration in tokay geckos. Finally, the results of our current study show that individual responses across tests were not repeatable (R = 0.037) while we found low individual repeatability in our first study (R = 0.120) [26]. In our current study, effects of prey species and habituation to novel space could be causing a decrease in individual repeatability. Further studies are, therefore, needed to either confirm that neophobia is not a single trait in the tokay gecko as we have discussed previously [26] or that it forms a single trait but it can be easily masked by test specific effects.

Finally, we found some effects of environmental variables on different measures of neophobia. The time to attack food decreased with rising temperature. Because temperature is related to digestion in ectotherms such as lizards [49], we would expect food motivation to be related to temperature. Furthermore, we found that space neophobia decreased with increasing time of day in the current study but this effect vanished when all three trials (both studies) were taken into account. Our geckos’ main activity period is in the morning and we aim to conduct testing during this time. However, this was not possible in the space neophobia test due to the trials being 20 minutes long. Consequently, we tested lizards in a different random order each trial to account for this effect of time of day. It seems with more trials conducted in a random order this effect, indeed, disappears as intended by randomising testing order.

## Conclusion

Our study demonstrates that the mere presence of a conspecific has no effect on neophobic responses in the tokay gecko, while pair association strength did affect space neophobia. Our results add important new insights into the influence of social context on neophobia in a facultative social lizard species. Our findings highlight that only considering presence/ absence of a conspecific in social species can lead to incomplete or possibly even false conclusions. We were also able to replicate our previous finding that tokay geckos hesitate to attack unfamiliar food and food near novel objects but hesitate the most when entering novel space. However, we also found that prey species, temperature, time of day and habituation are factors that need to be considered in future studies. Consequently, only object neophobia and exploration proofed repeatable, and therefore, reliable measures in tokay geckos. Our study brings us a step closer in better understanding how sociality influences cognition in lizards but already proved that social lizards are an excellent model to investigate this relationship.

## Data Availability

All data generated for this study are available at the Open Science Framework (OSF). Link for review purposes: https://osf.io/h8nzs/?view_only=b712aac13a404912b7be0413d9f04ea9 The code used for analysis is available at the Open Science Framework (OSF). Link for review purposes: https://osf.io/h8nzs/?view_only=b712aac13a404912b7be0413d9f04ea9

## Acknowledgements

We thank Lauriane Begue, Fabrice Bucheli, Annick Bühler and Laurin Ammon for their help with inter-observer reliability. We also thank the members of the Division of Behavioural Ecology of the University of Bern for their helpful comments on the first draft of this manuscript. This work was supported by the Austrian Science Fund (FWF) [grant P 31518, PI: ER] and the Swiss National Science Foundation (SNSF) [grant 197921, PI: ER].

## Author contributions

BS - Conceptualization; BS - Data curation; BS - Formal analysis; ER - Funding acquisition; BS - Investigation; BS - Methodology; BS, ER - Project administration; BS, ER - Resources; BS - Validation; BS - Visualization; BS, ER - Roles/Writing - original draft; BS, ER - Writing - review & editing.

## Supplementary figures

**Figure S1.**
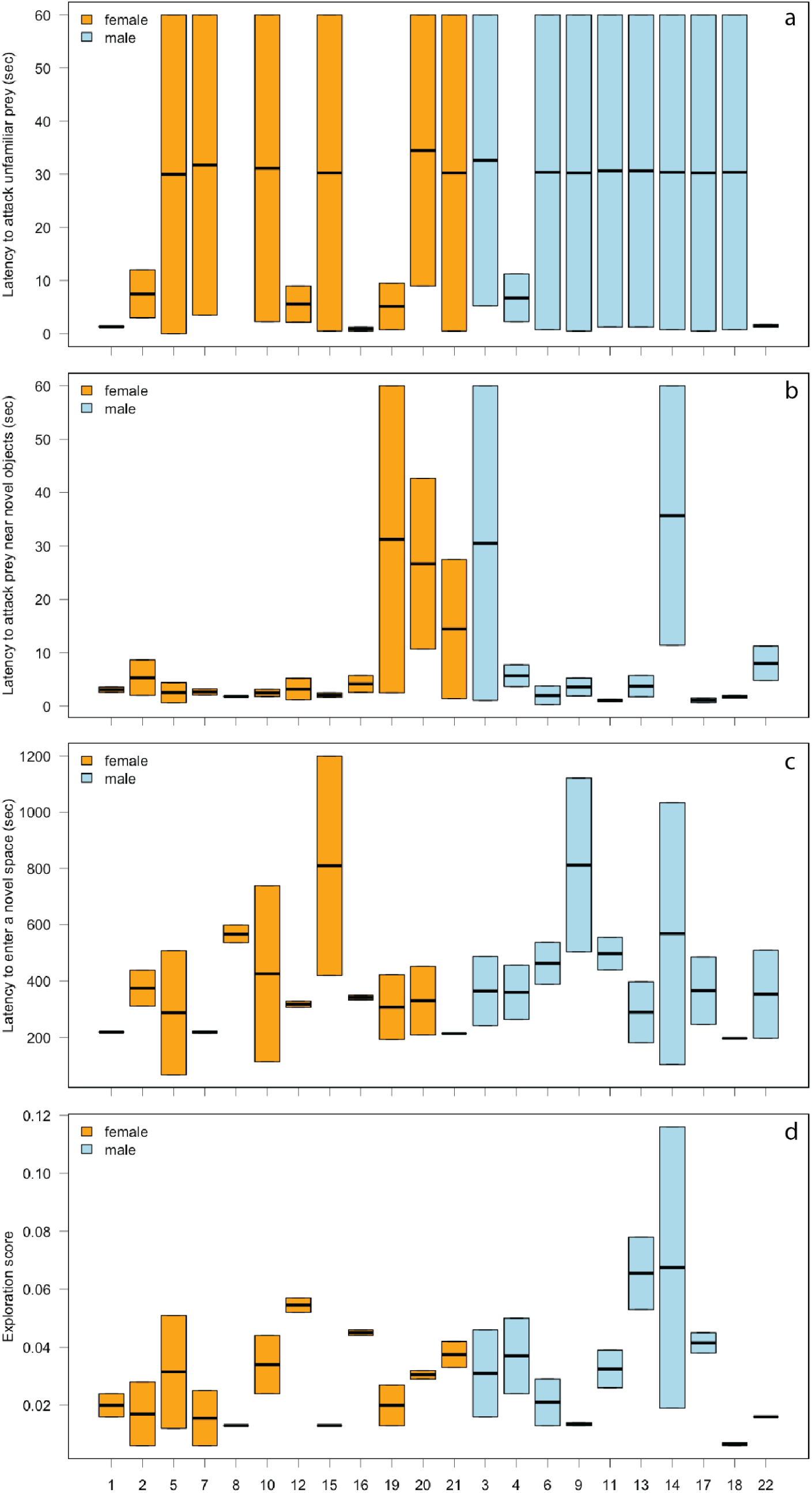
Individual responses towards novel stimuli and exploration in a novel space separated into females (orange) and males (blue) in the pair housing experiment. (a) Individual attack latencies towards unfamiliar prey in the food neophobia test. (b) Individual attack latencies towards familiar prey near novel objects in the object neophobia test. (c) Individual exit latencies into a novel space. (d) Individual exploration scores in a novel space. The bold line within boxes indicates the median, the upper box edges give the upper quartile and the lower box edges the lower quartile (outliers are not shown). Whiskers not visible because individuals were only tested in two trials.

## Supplementary tables

**Table S1.**
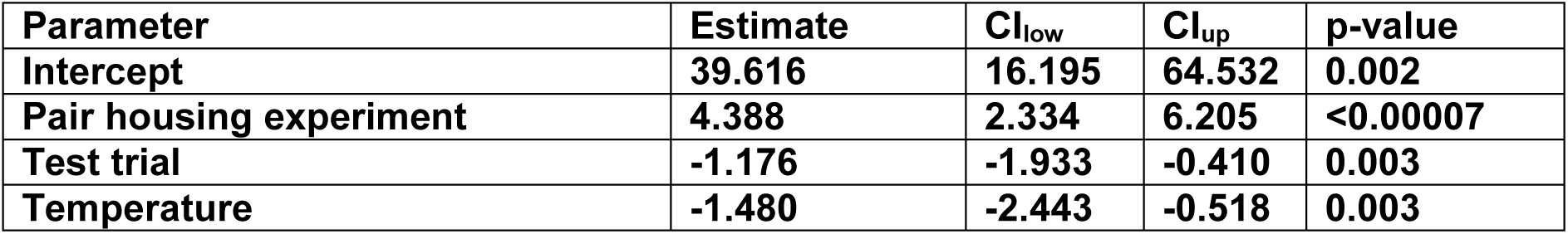
Estimates and tests statistics from the model comparing food neophobia across the two studies (single and pair housing). The model was run with 6000000 iterations, a burn in of 10000 and a thinning interval of 400. The response variable was log-transformed. CI_low_ – lower 95% Credible Interval, CI_up_ – upper 95% Credible Interval. Significant (p < 0.05) parameters are highlighted in bold.

**Table S2.** Estimates and tests statistics from the model comparing object neophobia across the two studies (single and pair housing). The model was run with 6000000 iterations, a burn in of 10000 and a thinning interval of 400. The response variable was log-transformed. CI_low_ – lower 95% Credible Interval, CI_up_ – upper 95% Credible Interval. Significant (p < 0.05) parameters are highlighted in bold.

| Parameter | Estimate | $CI_{low}$ | $CI_{up}$ | p-value |
| --- | --- | --- | --- | --- |
| Intercept | **1.738** | **0.947** | **2.495** | **<0.00007** |
| Pair housing experiment | 0.367 | -0.638 | 1.435 | 0.485 |
| Test trial | -0.283 | -0.765 | 0.204 | 0.244 |

**Table S3.** Estimates and tests statistics from the model comparing space neophobia across the two studies (single and pair housing). The model was run with 6000000 iterations, a burn in of 10000 and a thinning interval of 400. The response variable was log-transformed. CI_low_ – lower 95% Credible Interval, CI_up_ – upper 95% Credible Interval. Significant (p < 0.05) parameters are highlighted in bold.

| Parameter | Estimate | $CI_{low}$ | $CI_{up}$ | p-value |
| --- | --- | --- | --- | --- |
| Intercept | **7.025** | **6.469** | **7.605** | **<0.00007** |
| Pair housing experiment | 0.503 | -0.100 | 1.113 | 0.107 |
| Test trial | **-0.627** | **-0.982** | **-0.259** | **0.002** |
| Individual testing order | -0.025 | -0.075 | 0.026 | 0.340 |

**Table S4.** Estimates and tests statistics from the model looking at the relationship between food neophobia and pair association strength. The model was run with 10000000 iterations, a burn in of 10000 and a thinning interval of 400. The response variable was log-transformed. CI_low_ – lower 95% Credible Interval, CI_up_ – upper 95% Credible Interval. Significant (p < 0.05) parameters are highlighted in bold.

| Parameter | Estimate | $CI_{low}$ | $CI_{up}$ | p-value |
| --- | --- | --- | --- | --- |
| Intercept | **40.511** | **12.966** | **67.202** | **0.005** |
| Unfamiliar stimulus | **1.137** | **0.410** | **1.844** | **0.002** |
| Bond quality | 0.057 | -2.062 | 2.223 | 0.961 |
| Trial | **-1.362** | **-2.337** | **-0.398** | **0.007** |
| Temperature | **-1.530** | **-2.576** | **-0.471** | **0.006** |
| Unfamiliar stimulus – bond | 0.279 | -1.901 | 2.306 | 0.792 |
| Bond – trial | 1.453 | -1.369 | 4.210 | 0.294 |

**Table S5.**
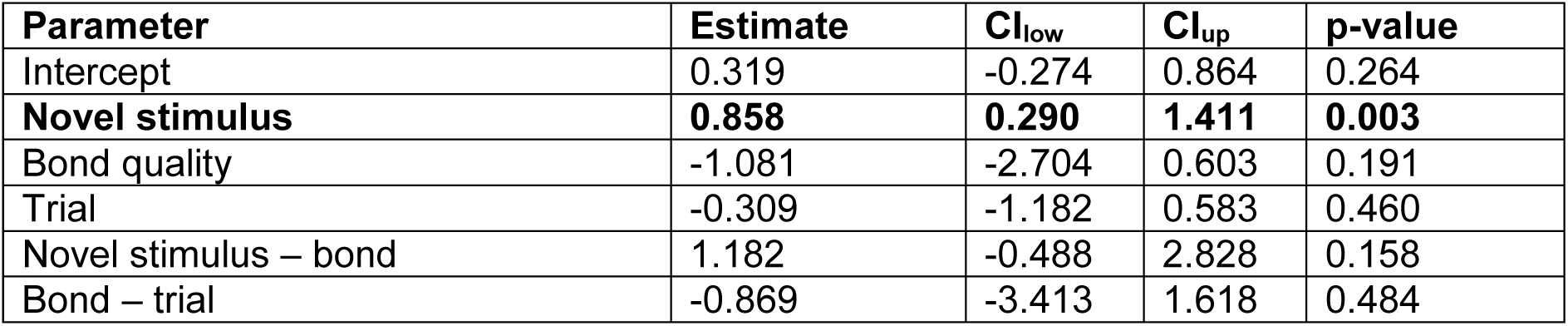
Estimates and tests statistics from the model looking at the relationship between object neophobia and pair association strength. The model was run with 10000000 iterations, a burn in of 10000 and a thinning interval of 400. The response variable was log-transformed. CI_low_ – lower 95% Credible Interval, CI_up_ – upper 95% Credible Interval. Significant (p < 0.05) parameters are highlighted in bold.

**Table S6.** Estimates and tests statistics from the model looking at the relationship between space neophobia and pair association strength. The model was run with 10000000 iterations, a burn in of 10000 and a thinning interval of 400. The response variable was log-transformed. We also present results from the PostHoc Estimated marginal means of linear trends. CI_low_ – lower 95% Credible Interval, CI_up_ – upper 95% Credible Interval. Significant (p < 0.05) parameters are highlighted in bold.

| Parameter | Estimate | $CI_{low}$ | $CI_{up}$ | p-value |
| --- | --- | --- | --- | --- |
| **Intercept** | **6.402** | **6.040** | **6.761** | **0.0005** |
| Bond quality | 0.101 | -0.579 | 0.742 | 0.761 |
| **Trial** | **-0.631** | **-0.945** | **-0.323** | **0.0002** |
| Individual testing order | -0.048 | -0.098 | 0.001 | 0.057 |
| **Bond – trial** | **1.050** | **0.109** | **1.942** | **0.027** |
| **PostHoc estimated marginal means** |  |  |  |  |
| **Trial** | **Trend of bond** | **$CI_{low}$** | **$CI_{up}$** |  |
| Trial 1 | 0.103 | -0.548 | 0.776 |  |
| Trial 2 | 1.150 | 0.389 | 1.904 |  |

**Table S7.** Estimates and tests statistics from the model looking at the relationship between exploration and pair association strength. The model was run with 6000000 iterations, a burn in of 10000 and a thinning interval of 400. The response variable was log-transformed. CI_low_ – lower 95% Credible Interval, CI_up_ – upper 95% Credible Interval. Significant (p < 0.05) parameters are highlighted in bold.

| Parameter | Estimate | $CI_{low}$ | $CI_{up}$ | p-value |
| --- | --- | --- | --- | --- |
| **Intercept** | **-3.900** | **-4.246** | **-3.554** | **<0.00007** |
| Bond quality | 0.475 | -0.568 | 1.463 | 0.338 |
| **Trial** | **0.471** | **0.023** | **0.917** | **0.040** |
| Bond quality – trial | -0.763 | -2.081 | 0.507 | 0.238 |

**Table S8.**
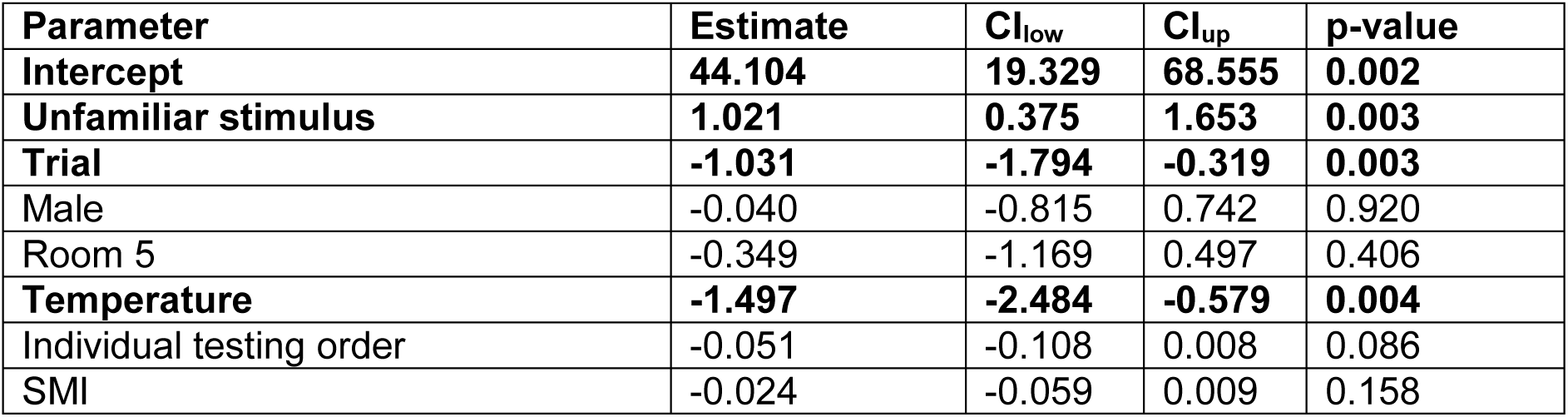
Estimates and tests statistics from the model looking at food neophobia. The model was run with 6000000 iterations, a burn in of 10000 and a thinning interval of 400. The response variable was log-transformed. CI_low_ – lower 95% Credible Interval, CI_up_ – upper 95% Credible Interval. Significant (p < 0.05) parameters are highlighted in bold. SMI – Scaled mass index (body condition).

| Parameter | Estimate | $CI_{low}$ | $CI_{up}$ | p-value |
| --- | --- | --- | --- | --- |
| **Intercept** | **44.104** | **19.329** | **68.555** | **0.002** |
| **Unfamiliar stimulus** | **1.021** | **0.375** | **1.653** | **0.003** |
| **Trial** | **-1.031** | **-1.794** | **-0.319** | **0.003** |
| Male | -0.040 | -0.815 | 0.742 | 0.920 |
| Room 5 | -0.349 | -1.169 | 0.497 | 0.406 |
| **Temperature** | **-1.497** | **-2.484** | **-0.579** | **0.004** |
| Individual testing order | -0.051 | -0.108 | 0.008 | 0.086 |
| SMI | -0.024 | -0.059 | 0.009 | 0.158 |

**Table S9.** Estimates and tests statistics from the model looking at responses (attack latency) towards familiar and unfamiliar rey types across both studies. The model was run with 6000000 iterations, a burn in of 10000 and a thinning interval of 400. The response variable was log-transformed. CI_low_ – lower 95% Credible Interval, CI_up_ – upper 95% Credible Interval. For contrasts we assumed statistical significance if the 95% credible interval did not cross 0. Significant (p < 0.05) parameters are highlighted in bold.

| Parameter | Estimate | $CI_{low}$ | $CI_{up}$ | p-value |
| --- | --- | --- | --- | --- |
| **Intercept** | **0.560** | **0.263** | **0.849** | **0.0001** |
| Mealworm | 0.435 | -0.167 | 1.020 | 0.151 |
| Locust | 0.271 | -0.312 | 0.830 | 0.350 |
| **Silk moth larvae** | **2.452** | **1.817** | **3.076** | **<0.00007** |
| Cockroach | -0.505 | -1.098 | 0.101 | 0.100 |
| **Contrasts** |  |  |  |  |
| Mealworm – locust | 0.164 | -0.588 | 0.895 |  |
| **Mealworm – silk moth larvae** | **-2.017** | **-2.800** | **-1.254** |  |
| **Mealworm – cockroach** | **0.940** | **0.170** | **1.701** |  |
| **Locust – silk moth larvae** | **-2.181** | **-2.944** | **-1.387** |  |
| **Locust – cockroach** | **0.776** | **0.025** | **1.521** |  |
| **Silk moth larvae – cockroach** | **2.957** | **2.108** | **3.727** |  |

**Table S10.** Estimates and tests statistics from the model looking at object neophobia. The model was run with 6000000 iterations, a burn in of 10000 and a thinning interval of 400. The response variable was log-transformed. CI_low_ – lower 95% Credible Interval, CI_up_ – upper 95% Credible Interval. Significant (p < 0.05) parameters are highlighted in bold. SMI – Scaled mass index (body condition).

| Parameter | Estimate | $CI_{low}$ | $CI_{up}$ | p-value |
| --- | --- | --- | --- | --- |
| Intercept | 5.091 | -18.314 | 28.505 | 0.673 |
| **Novel stimulus** | **0.992** | **0.462** | **1.517** | **0.003** |
| Trial | -0.143 | -0.840 | 0.517 | 0.688 |
| Male | 0.351 | -0.371 | 1.073 | 0.335 |
| Room 5 | 0.0.128 | -0.562 | 0.880 | 0.722 |
| Temperature | -0.095 | -0.970 | 0.811 | 0.841 |
| Individual testing order | -0.026 | -0.075 | 0.026 | 0.303 |
| SMI | -0.020 | -0.049 | 0.007 | 0.150 |

**Table S11.**
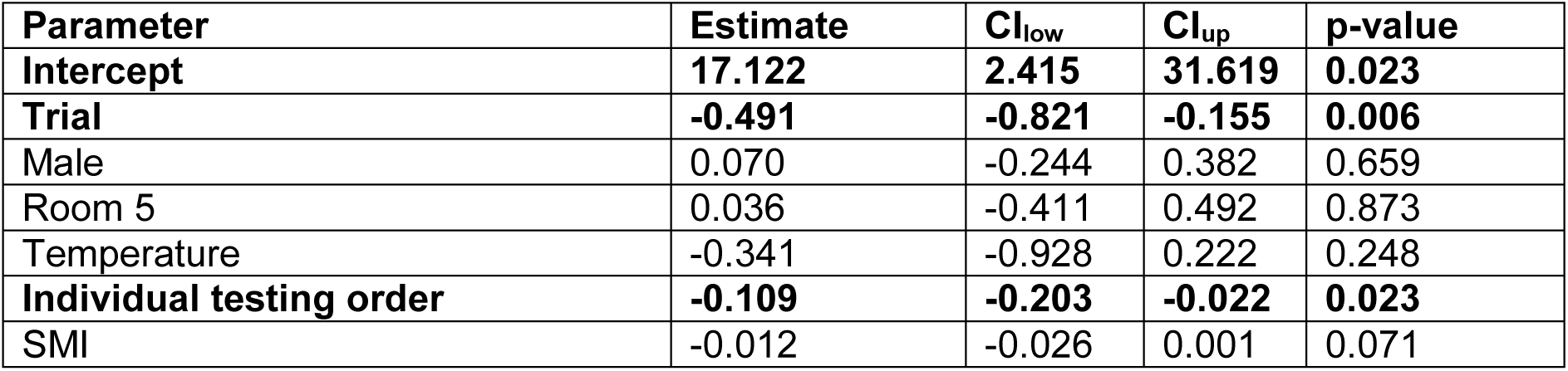
Estimates and tests statistics from the model looking at exit latency during the space neophobia. The model was run with 6000000 iterations, a burn in of 10000 and a thinning interval of 400. The response variable was log-transformed. CI_low_ – lower 95% Credible Interval, CI_up_ – upper 95% Credible Interval. Significant (p < 0.05) parameters are highlighted in bold. SMI – Scaled mass index (body condition).

**Table S12.** Estimates and tests statistics from the model looking at exploration during the space neophobia. The model was run with 16000000 iterations, a burn in of 10000 and a thinning interval of 400. The response variable was log-transformed. CI_low_ – lower 95% Credible Interval, CI_up_ – upper 95% Credible Interval. Significant (p < 0.05) parameters are highlighted in bold. SMI – Scaled mass index (body condition).

| Parameter | Estimate | $CI_{low}$ | $CI_{up}$ | p-value |
| --- | --- | --- | --- | --- |
| Intercept | -3.321 | -23.783 | 17.196 | 0.741 |
| **Trial** | **0.459** | **0.010** | **0.896** | **0.044** |
| Male | -0.030 | -0.426 | 0.526 | 0.902 |
| Room 5 | -0.002 | -0.648 | 0.647 | 0.992 |
| Temperature | 0.007 | -0.778 | 0.826 | 0.982 |
| Individual testing order | 0.036 | -0.092 | 0.164 | 0.574 |
| SMI | -0.014 | -0.034 | 0.006 | 0.155 |

**Table S13.** Estimates and tests statistics from the model looking at the difference in relative latency across neophobia tests. The model was run with 6000000 iterations, a burn in of 10000 and a thinning interval of 400. The response variable was log-transformed. CI_low_ – lower 95% Credible Interval, CI_up_ – upper 95% Credible Interval. Significant (p < 0.05) parameters are highlighted in bold.

| Parameter | Estimate | $CI_{low}$ | $CI_{up}$ | p-value |
| --- | --- | --- | --- | --- |
| **Intercept** | **-2.546** | **-3.060** | **-2.068** | **<0.00007** |
| Object neophobia test | -0.203 | -0.832 | 0.425 | 0.528 |
| **Space neophobia test** | **1.336** | **0.729** | **1.977** | **<0.00007** |
| **Contrasts** |  |  |  |  |
| **Object – space neophobia test** | **-1.539** | **-2.117** | **-0.900** |  |

**Table S14.** Results from the Spearman rank correlation tests looking at correlations of neophobic responses across tests within the pair housing experiment.

| Contrast | Correlation coefficient | S-value | p |
| --- | --- | --- | --- |
| Food – object neophobia | -0.188 | 1829.1 | 0.415 |
| Food – space neophobia | 0.161 | 1292.8 | 0.487 |
| Object – space neophobia | -0.144 | 2015.1 | 0.524 |

